# Comparative analysis of metabarcoding and metagenomics for fish biodiversity estimates using standard and novel high-flow filtration methods

**DOI:** 10.64898/2025.12.15.694277

**Authors:** Daniël van Berkel, Niels W. P. Brevé, Mark de Boer, Emmanuel G. Reynaud, Reindert Nijland

## Abstract

Molecular techniques involving environmental DNA (eDNA) are increasingly used for aquatic species detection. Metabarcoding, a widely adapted technique, suffers from primer bias: uneven amplification of species due to primer mismatches. The primer bias can be eliminated by omitting PCR, thereby sequencing all eDNA in a sample. This method, known as metagenomics, offers potential benefits for relative abundance estimates and epigenetic modifications, but is seldom applied to eukaryotic communities and eDNA.

This study uses an expanded two-by-two design to compare fish species detection between multi-marker metabarcoding and metagenomics using two filter types (conventional versus high-flow). Environmental DNA was collected in a controlled setup and two field settings, which contained several fish species including European sturgeon (*Acipenser sturio*). Moreover, we explore methylation patterns obtained from nanopore native sequencing.

All species present in the controlled environment were detected using both metabarcoding and metagenomics. In field settings, metagenomics detected more species than metabarcoding. High-flow filters recovered more species across all sequencing datasets, except in metabarcoding of field settings. Relative read counts between metabarcoding and metagenomics illustrate primer bias is present in the used primer sets. Most fish metagenomic sequences were identified as *A. sturio* across all eDNA samples. We observed three base modifications on the 18S region of *A. sturio*, where three sites showed different methylation patterns between eDNA samples.

Our results demonstrate that metabarcoding and metagenomics function complementary in species detection and metagenomics provides additional insights into base modifications. Moreover, high-flow filters offer strong potential for improved species detection in various environments.

## Introduction

Molecular techniques involving environmental DNA (eDNA) are used increasingly for aquatic species detection and are, in some cases, able to replace traditional monitoring methods (Takahashi et al. 2023). One of the most widely used molecular techniques is DNA metabarcoding, where several species or taxa of interest are amplified with universal primers, thereby providing insight into the species composition and corresponding biodiversity of certain taxonomic groups (Taberlet et al. 2012, Deiner et al. 2017b). Metabarcoding of eDNA is relatively easy, cheap, does not require expert taxonomists and the method has provided similar results to traditional visual identification methods (e.g. pulse fishing, bottom trawling, BRUVs) (Olds et al. 2016, Liu et al. 2022, Thompson et al. 2024). However, certain limitations remain with this molecular technology.

One of the disadvantages of eDNA metabarcoding is the abundance estimation of species; relative read counts following PCR amplification cannot be translated to relative abundance due to several biases. The most prominent bias which occurs after eDNA collection is primer bias, which occurs due to mismatches in the primer binding sites (Polz and Cavanaugh 1998). Universal primer sets are an effective tool for amplifying several target species and are therefore commonly used in biodiversity assessments (e.g. Deiner et al. 2017a, Macé et al. 2024, Maggini et al. 2024, Doorenspleet et al. 2025). However, universal primer sets rarely amplify all target species equally due to mismatches in primer binding sites, which create a severe bias towards the relative read count of certain species (Piñol et al. 2015). This primer bias is evident within most taxonomic groups, including elasmobranchs (Elasmobranchii) and ray-finned fish (Actinopterygii) (Zhang et al. 2020, West et al. 2024).

The primer bias can be eliminated by omitting the PCR amplification step altogether, thereby sequencing all eDNA present in a water sample. This technique is known as metagenomics, or sometimes ‘shotgun sequencing’ or ‘native sequencing’, depending on the sequencing technique provider (Zepeda Mendoza et al. 2015, Quince et al. 2017, Liu et al. 2025). Besides the more accurate abundance estimates, metagenomics sequencing data could also provide (mito)genome assemblies or genome skimming (depending on sequencing depth) and epigenetic modifications, depending on the abundance of the target species’ DNA and its sequencing coverage (Zhao et al. 2023, Lu et al. 2025). This additional information allows for deeper insights in biodiversity, not only looking into species diversity, but also genetic diversity and life-history differences. Metagenomics has already been used for three decades to study aquatic prokaryotic communities (Stein et al. 1996) and has recently also been applied to eukaryotes in aquatic environments. The technique has been used in eDNA for various taxonomic groups, but the main focus within marine eukaryotes is on fish, as these generally have sufficient genomic reference databases to identify taxa to species level (Hotaling et al. 2021).

Despite the advantages and increasing interest in applying metagenomics to eDNA fish communities, its use can be challenging due to the low abundance of fish eDNA in a water sample. Few studies report the exact proportion of fish DNA in aquatic metagenomic eDNA, but generally 1% of all eDNA reads are classified as chordates (Cowart et al. 2018, Lui and Nielsen 2024, Curto et al. 2025, Patin et al. 2025). This low proportion means deep sequencing is essential to obtain sufficient sequencing depth for thorough fish biodiversity estimates. Deep sequencing is more costly due to the amount of sequencing data that is collected. Since large DNA quantities required as input, multiple water replicates are often collected to obtain sufficient DNA quantities and simultaneously reduce variance between water samples. Fish biodiversity estimates could be improved by sequencing deeper or by increasing the proportion of fish DNA collected on an eDNA filter. As fish regularly reside in murky or sandy environments, the filtering capacities of the widely used 0.45 μm and 1.2 μm disk filters are often limited by suspended sediment and micro-organisms, thereby limiting target DNA yield. Using a coarse (greater pore size), high-flow, filter for eDNA filtration could improve filtering and on-target metagenomic sequencing efficiency by allowing smaller fragments to pass through, while retaining clumps of cells from multicellular (fish) origin. By doing so, the filtration volume can be increased (yielding higher target DNA quantities) while simultaneously decreasing the proportion of non-target DNA in the filter (Nester et al. 2024). Additionally, the increased filtration volume could omit the need for field replicates, otherwise necessitated by low filtration volumes, reducing the costs and time spend during fieldwork (Cantera et al. 2019, How et al. 2024). While the use of coarser filters and increased filtering volumes has been shown to improve species detection and abundance estimates in metabarcoding (Govindarajan et al. 2022, Nester et al. 2024), it is unknown if these adaptations provide the same benefits to metagenomic fish detection.

In this study, we compare fish species detection using multi-marker metabarcoding and metagenomics of two eDNA filter types: a widely implemented 1.2 μm disk filter and alternatively, a coarse, high-flow filter. We collected water from a controlled environment and from two field settings, and then compare the fish species observed with metabarcoding and metagenomics using both filter types. In addition, we explore a potential benefit of metagenomics: epigenetic base modifications. We expect the following four outcomes: a) less species will be detected using metagenomics compared to metabarcoding, as some reference genomes of target species are not publicly available and some rare species could go undetected due to extremely low DNA fragment numbers, b) metagenomics will provide different relative read counts due to the lack of primer bias, c) the increased throughput of the high-flow filters will increase the proportion of fish DNA collected, increasing on-target metagenomic sequencing efficiency, and d) methylation patterns will differ between animals from varying environments and different life stages.

## Methods

### Water collection and filter types

Water was collected from both a controlled environment (Diergaarde Blijdorp/Rotterdam Zoo, the Netherlands) and two field settings (Biesbosch nature reserve), both with a focus on European sturgeon (*Acipenser sturio*) because of the ecological interest in the species (Fig. 1). At Diergaarde Blijdorp, water was taken directly from an aquarium containing three 14-year-old European sturgeons (ca. 140 cm) and six other fish species represented by nine individuals: four *Dasyatis tortonesei* individuals (two disk size 80-100 cm, two disk size 30 cm), one *Labrus mixtus* (length 30 cm), one *Mustelus asterias* (length 100 cm), one *Raja brachyura* (disk size 50 cm), one *Scomber scombrus* (length 40 cm), and one S*parus aurata* (length 45 cm). At the Biesbosch, a Dutch freshwater nature reserve, 29 juvenile (16 month old, ca. 25 cm) European sturgeons were reintroduced during an event on June 2^nd^, 2023 (Fig. 1). Here, water was collected from a storage vessel (samples labeled: Biesbosch Holding tank) where the sturgeons had been kept for a week before being reintroduced (51°45’56”N, 4°46’14”E). At both locations, water was filtered directly after collection using three Sartorius 1.2 μm 45 mm disk filters (Göttingen, Germany) (hereafter: conventional filters) and three Smoking cigarette filters (Ø 8mm, length 22 mm) (Barcelona, Spain), which have an approximate pore size of 3-6 μm (Idriss et al. 2024), allowing more flow (hereafter: high-flow filters). Two liters of water were filtered per filter. At the Biesbosch, three additional water samples were collected to determine the maximum filtration volume of the high-flow filters and to provide additional sequencing data on an individual level (Fig. 1). The first water sample was collected from the sturgeon’s store vessel where the water was filtered until the high-flow filter clogged, thereby obtaining the maximum filter volume. The other two water samples were collected from a bucket where a single juvenile European sturgeon was kept for several minutes, providing a capture-release scenario (samples labeled: Biesbosch Bucket). These two water samples provide insights on metagenomic sequencing data on an individual level rather than the population level of the zoo aquarium or sturgeon’s storage vessel. For both Biesbosch Bucket samples, two liters of water were filtered, one through a conventional filter, the other through a high-flow filter (Fig. 1).

**Figure 1.**
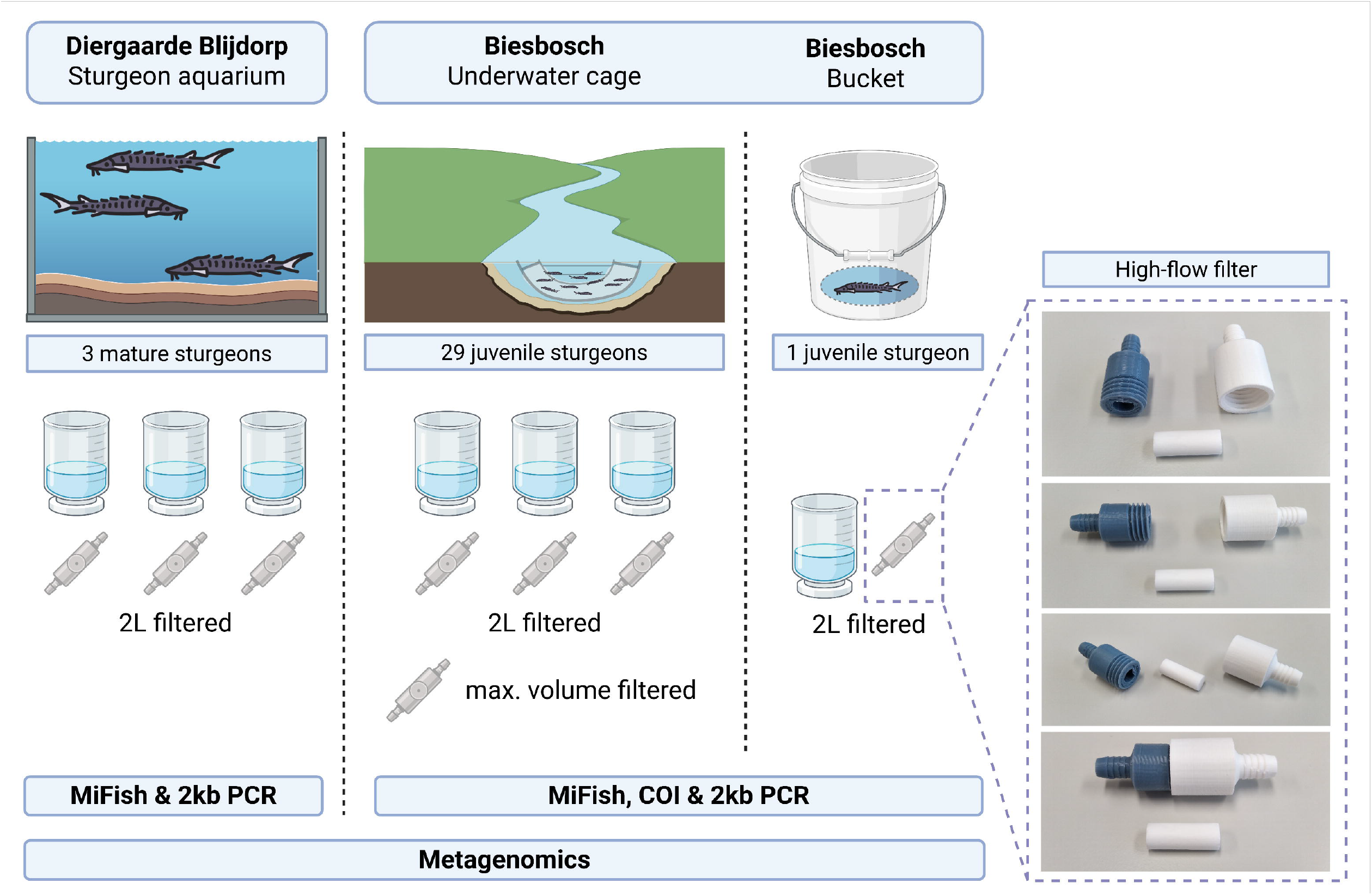
Graphical overview of water eDNA samples collected for this study.

At Blijdorp as well as Biesbosch, the filter setup for conventional filters consisted of three Rocker filter holders and 300 mL funnels (Kaohsiung, Taiwan), with individual collection vessels. The filter setup of the high-flow filters consisted of three high-flow filters placed in 3D printed filter holders (Saggiomo and Reynaud 2025) and a single collection vessel. All water was filtered using a Youcheng Zhixin VN-C4 pump (pumping rate up to 32 L/min, vacuum up to −90 kPa) (Bacheng town, China). Following water filtration, filters were directly placed in a 2 mL screw cap tubes pre-filled with 400 uL Zymo DNA/RNA Shield (Irvine, CA, USA) and placed on ice for short-term storage during transport. After transportation to the molecular laboratory, the tubes were stored at −20 °C until DNA isolation.

### eDNA extraction, metabarcoding and metagenomics

DNA was extracted from the filters using the Qiagen Gentra kit (Hilden, Germany). The DNA/RNA Shield was transferred from each screw cap to a 1.5 mL Eppendorf tube, transferring as much supernatant as possible. As the high-flow filters held more than 500 uL supernatant, it was split between two 1.5 mL Eppendorf tubes, yielding duplicate DNA isolations. One negative DNA control was included, consisting of 400 uL DNA/RNA shield. Then, Proteinase K was added according to the Gentra kit protocol. Samples were incubated for 2 hours at 55 °C, 500 rpm to lyse any tissue. The volume of protein precipitation buffer and isopropanol were adjusted to appropriate ratios. DNA was precipitated according to the Gentra protocol. Finally, the volume of DNA Hydration Solution was reduced to 56 uL to increase DNA concentrations and improve metagenomic sequencing. DNA was dissolved at 4 °C overnight, after which the quality and quantity were measured using the Thermo Scientific Nanodrop 1000 spectrophotometer (Waltham, MA, USA) and Invitrogen Qubit 2.0 fluorometer (Waltham, MA, USA).

For metabarcoding, three mitochondrial genome regions were amplified for all DNA isolations. The MiFish and 2kb primer sets were used to target fish and other vertebrates (Miya et al. 2015, Doorenspleet et al. 2025), whereas the Leray COI primer set was used to target metazoans (Leray et al. 2013). The COI and 2kb primer sets had ONT barcoding tails attached to ease the second-round barcoding PCR. In contrast, tagged primers were used for the MiFish PCR, which used unique 13 bp tags included in the forward and reverse primer, thereby omitting the need for a second round barcoding PCR (Srivathsan et al. 2019). The annealing section of the MiFish primer set consisted of the following forward and reverse primer, respectively: GTYGGTAAAWCTCGTGCCAGC & ATAGTGGGGTATCTAATCCYAGTTTG, the annealing section of the 2kb primer set consisted of the following forward and reverse primers, respectively: GGATTAGATACCCYACTATGC & GATTGCGCTGTTATCCCTAG, whereas the annealing section of the COI primer set consisted of the following forward and reverse primers, respectively: GGWACWGGWTGAACWGTWTAYCCYCC & TANACYTCNGGRTGNCCRAARAAYCA (Geller et al. 2013, Miya et al. 2015, Doorenspleet et al. 2021, 2025). Each DNA isolation (including the negative control) was amplified with triplicate PCR reactions where each reaction consisted of 5 μL Thermo Scientific Phire Tissue Direct Mastermix (Waltham, MA, USA), 0.2 μL primer mix (10mM forward + reverse), 0.5 μL DNA template and 4.3 μL nuclease free water. Three negative PCR controls were included in each PCR, where the DNA template was substituted for nuclease free water. The PCRs were performed in a Bio-Rad T100 (Hercules, CA, USA) where programs were specific for each primer set; all started with initial denaturation for 3 minutes at 98 °C, followed by 35 cycles of 10 seconds at 98 °C, 10 seconds at 59/57/55 °C (MiFish/2kb/COI), then 10/40 seconds at 72 °C (MiFish&COI/2kb) and lastly, a final elongation for 3 minutes at 72 °C. A 1.5 μL subsample of each PCR reaction was loaded on a 1% agarose gel and subsequently assessed using the BioRad GelDoc XR (Hercules, CA, USA). If successful, the COI and 2kb PCR replicates were pooled and a second-round barcoding PCR was initiated according to the Oxford Nanopore Technologies (ONT, Oxford, UK) EXP-PBC096 protocol. A 1.5 μL subsample of the barcoding PCR was again loaded on a 1% agarose gel and assessed using the Bio-Rad Geldoc XR. The tagged MiFish and barcoded COI and 2kb amplicons were pooled in equimolar ratios according to the amplicon length and brightness of the target band on gel. The end-prep and adapter ligation were performed according to the ONT SQK-LSK114 protocol. Sequencing was performed on a ONT R10.4.1 MinION flowcell using a ONT Mk1C device.

For metagenomics, the ONT Native barcoding protocol (SQK-NBD114.24) was used to sequence all DNA isolations. In case the DNA concentration of a DNA isolation was insufficient according to the protocol (i.e. 400 ng in 10 uL), the volume was reduced to approximately 10 μL using a Thermo Scientific SpeedVac DNA 130 (Waltham, MA, USA). If the total weight of a DNA isolation was insufficient (<400 ng), DNA isolation replicates were pooled and then concentrated. In case the total weight of a DNA isolation was less than 100 ng, 1 uL of DNA control strand (DCS, 10 ng) was added to improve the library preparation and subsequent sequencing. Sequencing was performed on a ONT 10.4.1 PromethION flowcell placed in a PromethION 24 device. All raw sequencing data is available on ENA under the project accession PRJEB96019.

### Species identification

Following sequencing, the raw metabarcoding data (POD5 files) were SUP-basecalled, trimmed and demultiplexed with dorado (version 0.7.0). As MiFish had tagged primers, these were identified during basecalling using the following command: dorado basecaller dna_r10.4.1_e8.2_400bps_sup@v4.3.0 /input_folder/ -r --no-trim --barcode-sequences tags. fa --barcode-arrangement tags.toml --min-qscore 10 > no_trim.bam. Adapters were then trimmed using the following command: dorado trim -- no-trim-primers no_trim.bam > trimmed.bam and data was then demultiplexed using: dorado demux trimmed.bam -- no-classify --no-trim --emit-fastq --output-dir /Demux_MiFish/. For COI and 2kb, data was basecalled using the following commands: dorado basecaller dna_r10.4.1_e8.2_400bps_sup@v4.3.0 /input_folder/ -r --trim adapters --min-qscore 10 > basecalled_COI_2kb.bam and then demultiplexed using: dorado demux basecalled_COI_2kb.bam --kit-name EXP-PBC096 --barcode-both-ends --emit-fastq -- output-dir /Demux_COI_2kb/. Raw metagenomic data (POD5 files) were also SUP-basecalled using dorado (version 0.7.0) using the following command: dorado basecaller dna_r10.4.1_e8.2_400bps_sup@v4.3.0 -r --kit-name SQK-NBD114-24 /input_folder/ --min-qscore 10 > data.bam and then demultiplexed using: dorado demux data.bam --kit-name SQK-NBD114-24 --barcode-both-ends. Metabarcoding sequences were processed using DECONA (https://github.com/saskia-oosterbroek/decona), which incorporates primer trimming, clustering, polishing and species identification using BLAST, with the following settings for MiFish: decona -i /Decona_MiFish -f -q 10 -l 150 -m 240 -g “GTYGGTAAAWCTCGTGCCAGC;max_error_rate=0.1;min_overlap=17…CAAACTRGGA TTAGATACCCCACTAT;max_error_rate=0.1;min_overlap=21” -c 0.96 -n 10 -r -o 1 -R 500 -k 6 -M -b /local/nt_euk, for COI: decona -i /Decona_COI -f -q 10 -l 300 -m 360 -g “GGWACWGGWTGAACWGTWTAYCCYCC;max_error_rate=0.1;min_overlap=20” -a “TGRTTYTTYGGNCAYCCNGARGTNTA;max_error_rate=0.1;min_overlap=22” -c 0.95 -n 10 -r -o 0.98 -R 500 -k 6 -M -b /local/nt_euk and for 2kb: decona -i /Decona_2kb -f -q 10 -l 1800 -m 2350 -g “GGATTAGATACCCYACTATGC;max_error_rate=0.1;min_overlap=17…CTAGGGATAAC AGCGCAATC;max_error_rate=0.1;min_overlap=17” -c 0.93 -n 10 -r -o 0.99 -R 500 -k 6 -M -b /local/nt_euk. Next, filtering steps were performed in Rstudio (version 2025.05.0) using R (version 4.5.1) and the dplyr package; reads were discarded if they were classified to family level or higher, had < 99 % percentage identity or had < 0.8 coverage (alignment length/amplicon length) (Wickham et al. 2014, R Core Team 2025).

For the metagenomic sequence data, a pre-selection using Kraken2 was performed to extract all potential fish reads, after which reads were identified to species level using BLAST (Wood et al. 2019, Sayers et al. 2025). First, a species list of fish occurring in Dutch waters was obtained from Naturalis (https://www.nederlandsesoorten.nl/content/mariene-soortenlijst). Species present in the Blijdorp aquarium or detected with metabarcoding were added to the species list. The scientific names of these species were then used as query in ncbi-datasets, through which reference genomes and mitochondrial genomes (mitogenomes) were obtained, if available (O’Leary et al. 2024). These reference (mito)genomes were then used to construct a Kraken2 reference database, where all metagenomics reads were processed with the following command: kraken2 fish_reference_genomes reads.fasta --confidence 0.25 output.txt > report.txt (Wood et al. 2019). The reads classified by Kraken2 were subsequently identified to species level using a local BLAST search with the nt_euk database (downloaded 22-8-2025) (Sayers et al. 2025) using the following command: blastn -max_target_seqs 15 - max_hsps 5 -dust yes -outfmt “6 qseqid pident length mismatch gapopen qstart qend sstart send evalue bitscore qlen slen sseqid salltitles sallseqid” -query kraken2_classified_reads_2.fasta -db NCBI_nt_euk_20250822/nt_euk -out BLAST_results.txt.

An additional Kraken2 database and BLAST database were constructed using the reference genome of *Acipenser sturio* (ENA accession: GCA_977009215), as this genome was not present in NCBI’s nt_euk database. All metagenomic reads were processed using the *A. sturio* Kraken2 reference database using a confidence score of 0.25 and subsequently processed by the *A. sturio* BLAST database using the same settings as the nt_euk BLAST. The output of the nt_euk BLAST and *A. sturio* BLAST were then merged, and a “best-hit” was obtained for each read in R (version 4.5.1) and Rstudio (version 25.05.0) using the dplyr package by selecting the species with the lowest E-value (Wickham et al. 2014, R Core Team 2025). In case multiple species had equal E-values, the lowest common taxa was selected as “species” taxa (e.g. *Acipenser* sp. or Acipenseridae sp.) After selecting the best hit, reads were selected with a minimum percentage identity of 99%, minimum query length of 150 bp, a minimum coverage (alignment length/query length) of 0.8 and identification to genus- or species level. In addition, species detections were considered invalid if less than 10 reads occurred in the entire dataset, if the proportion of that species within a single barcode was less than 0.01% or if less than three reads of that species occurred within a single barcode.

A graphical overview of collected eDNA samples was created with BioRender.com and Flaticon.com. A Venn diagram of species observed in each environment with each filter type and metabarcoding/-genomics was made with DeepVenn (Hulsen 2022). Figures indicating the proportion of fish and *Acipenser sturio* observed in metabarcoding and metagenomics were made using Microsoft Excel version 2502.

### Methylation calling

Methylation patterns of three different base modifications (5mC, 5mCG and 6mA) were analyzed for all eDNA samples from the metagenomic sequencing data. The reads identified as *Acipenser* spp. by the Kraken2 and BLAST pipeline were selected from the raw .pod5 files using pod5 (version 0.3.28), after which 5mC, 5mCG and 6mA methylation were called using dorado (version 0.7.0) and the following modified basecall models: dna_r10.4.1_e8.2_400bps_sup@v4.3.0_(5mC_5hmC@v1)(5mCG_5hmCG@v1)(6mA@v2). These reads were then mapped to the 18S gene of *Acipenser sturio* (NCBI accession number: AY544133) using the aligner function of dorado. Mapped reads with a minimum mapping quality of 60 and primary mapping were then selected, sorted and indexed into a new .bam file using samtools 1.17 (Danecek et al. 2021). Samples were clustered based on environment (Blijdorp, Biesbosch Holding tank & Biesbosch Bucket) and filter methods (conventional filter, high-flow filter and high-flow filter maximum volume). Then, modkit (version 0.2.1) was used to calculate coverage and average methylation per site, per group (https://github.com/nanoporetech/modkit). The modkit .bed output was transformed to a .txt file and used as input for methylKit (version 1.34.0) in R (version 4.5.1) (Akalin et al. 2012). The methylation patterns of ecological samples (Blijdorp, Biesbosch Holding tank, Biesbosch bucket and) and filter types (conventional filter and high-flow filter) were compared using the methylKit’s package using a minimum coverage of 3 times. Average coverage of methylated sites and average methylation rates were calculated using dplyr.

## Results

### Sequencing metabarcoding and metagenomics

For metabarcoding, the total read counts for the MiFish, COI and 2kb primer set after demultiplexing were 2,397,619, 505,360 and 136,098, respectively. After quality filtering and selecting target taxa, 1,519,676 (63.4%), 40,140 (7,94%) and 97,321 (71.5%) remained for the MiFish, COI and 2kb primer set, respectively. While effort was made to amplify all eDNA samples with all three primer sets, some eDNA samples did not amplify with the COI primer set or 2kb primer set. For the eDNA isolations with successful amplification, the average fish read count was 48,177 ± 56,008 (median ± IQR) for MiFish, 1,141 ± 1,702 (median ± IQR) for COI and 6,420 ± 4,430 (median ± IQR) for 2kb.

For metagenomics, 23,386,873 reads and 39.18 Gb were sequenced in total. Following demultiplexing, removal of the DCS, quality filtering and selection of target taxa, the minimum read length was 150 bp and the maximum read length 28,023 bp. eDNA isolations consisted of 51,303 ± 118,764 (median ± IQR) reads and 5,039,533 ± 10,096,765 (median ± IQR) bp. The mean read length was highest in Blijdorp (586 bp) and lowest in Biesbosch Holding tank eDNA samples (414 bp). The mean fragment length of the high-flow filters was 10% lower than conventional filters in the Biesbosch Bucket samples (465 bp versus 523 bp), but 30% higher in Blijdorp samples (605 bp versus 462 bp) samples and even 60% higher in the Biesbosch Holding tank samples (474 bp versus 297 bp) (Supplementary Table 1).

All amplicon sequencing described used approximately 25% of a MinION flowcell’s full capacity, meaning that metabarcoding sequencing costs (excluding PCR and library prep) were less than €12 per eDNA sample. Metagenomic sequencing required a full PromethION flowcell, leading to sequencing costs of €60 per eDNA sample (excluding library prep).

### Species richness

Metabarcoding analysis resulted in 15 species detected in Blijdorp eDNA, of which all fifteen were observed with the high-flow filters and seven with the conventional filter (Fig. 2).

Similarly, metabarcoding analysis detected five fish species in the Biesbosch Holding tank, three of which were observed with the high-flow filters and four with the conventional filter. Metagenomic analyses resulted in eight species detected in Blijdorp eDNA and ten species detected in Biesbosch eDNA (Fig. 2). Using metagenomics, more species were detected using the high-flow filter than the conventional filter in Blijdorp eDNA (8 versus 5) as well as in Biesbosch eDNA (10 versus 8). In Blijdorp eDNA, one species (*Sprattus sprattus*) was not detected with metabarcoding, despite the detection with metagenomics. The opposite was observed in Biesbosch eDNA samples, where one species (*Neogobius fluviatilis*) was detected with metabarcoding, but not with metagenomics (Fig. 2).

**Figure 2.**
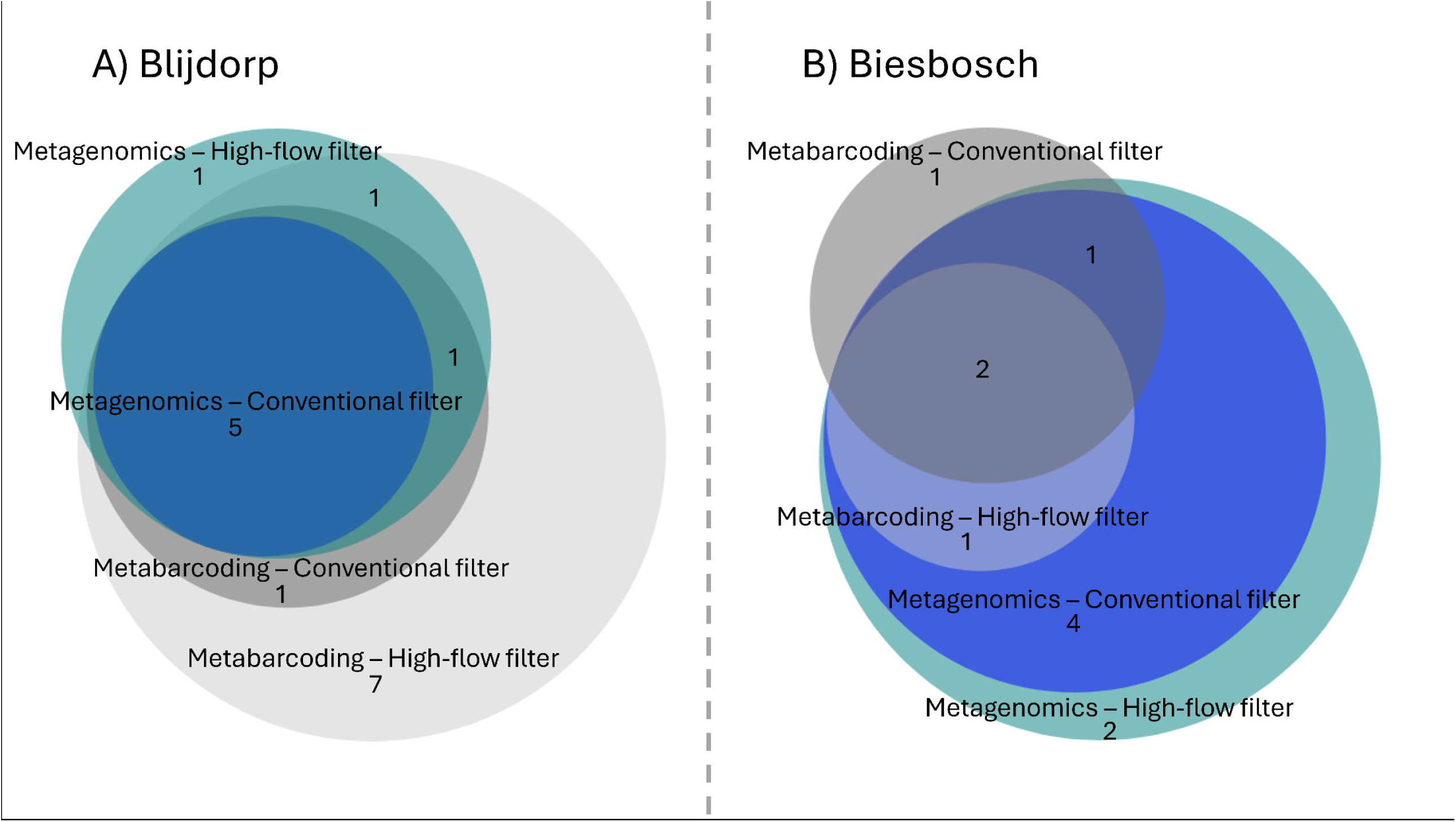
Venn-diagram of species detected two locations, using metabarcoding and metagenomics, and using two filter types; Venn-diagram of species detected at A) zoo aquarium and B) sturgeon release, using multi-marker metabarcoding and metagenomics, and using two filter types: conventional 1.2 μm disk filter and a high-flow cigarette filter. Panels A and B are not to scale to each other.

### Relative read counts

In Blijdorp, differences were observed in relative read counts of metabarcoding and metagenomics of the seven species that were detected with both methods. The seven species observed in MiFish metabarcoding and the metagenomics datasets were: *Acipenser sturio, Dasyatis tortonesei, Labrus mixtus, Mustelus asterias, Raja brachyura, Scomber scombrus* and *Sparus aurata. Acipenser sturio* was the most dominant species in metabarcoding as well as metagenomics, contributing on average 72.5% to all MiFish fish reads and 91.1% to all metagenomic fish reads. A similar pattern was observed for *Raja brachyura*, where the contribution to the total fish read count was higher in metagenomics (6.83%) than in metabarcoding (1.20%). In the other five overlapping species, a higher relative read count was observed in metabarcoding than in metagenomics. The difference in contribution to total fish read count between metabarcoding and metagenomics ranged between a two-fold difference for *M. asterias* (0.87% in metabarcoding, 0.43% in metagenomics) to a 400-fold difference in *D. tortonesei* (11.0% in metabarcoding, 0.026% in metagenomics) (Supplementary Table 2 & Supplementary Table 3).

The metagenomic relative read counts showed varying correlation with the abundance and biomass of fish present in the Blijdorp aquarium. The species with the most biomass was *Acipenser sturio*, with three mature individuals. This species also contributed to the majority of metagenomic fish reads. While two adult *Dasyatis tortonesei* (disk size 80-100 cm) and two juvenile (disk size 30 cm) were also present, only 0.026% of metagenomic detected in a single eDNA sample were attributed to this species. The other ray species, *Raja brachyura*, was represented by a single individual (disk size 50 cm) and contributed on average 6.83% to metagenomic fish reads. The last elasmobranch, a single mature *Mustelus asterias*, contributed only 0.43% to the metagenomic fish read count. The three teleosts did not contribute consistently to the fish metagenomic read count. The single S*parus aurata* (length 45 cm) contributed on average 0.34%, the single *Scomber scombrus* (length 40 cm) contributed on average 0.99% and the single *Labrus mixtus* (length 30 cm) contributed only 0.12% in a single eDNA sample (Supplementary Table 4).

### Fish classification rates

Filter type did not affect the percentage of MiFish reads classified as fish (hereafter: (fish) classification rates) for Blijdorp eDNA nor Biesbosch eDNA. However, classification rates were higher for Blijdorp eDNA than for Biesbosch eDNA with on average 72.5% and 54.2% classified, respectively. Increasing the water volume filtered with the high-flow filter increased the fish classification rate in the COI and 2kb primer set, but not in the MiFish primer set (Fig. 3). The vast majority of metabarcoding reads was identified as *Acipenser sturio* for the controlled- and field setting and both filter types, with 56% to 93% off all fish reads classified as *A. sturio* (Supplementary Table 2). In addition to the properly matching *Acipenser sturio* amplicons with a percentage identity of >99%, amplicons were observed in the metabarcoding data across all eDNA samples and across all three primer sets, which classified as other sturgeon species with low percentage identity scores (<97%). While these amplicons were discarded from the final dataset due to their low percentage identity, their origin is of interest due to their high relative read counts and are thus further elaborated in the Discussion section.

**Figure 3.**
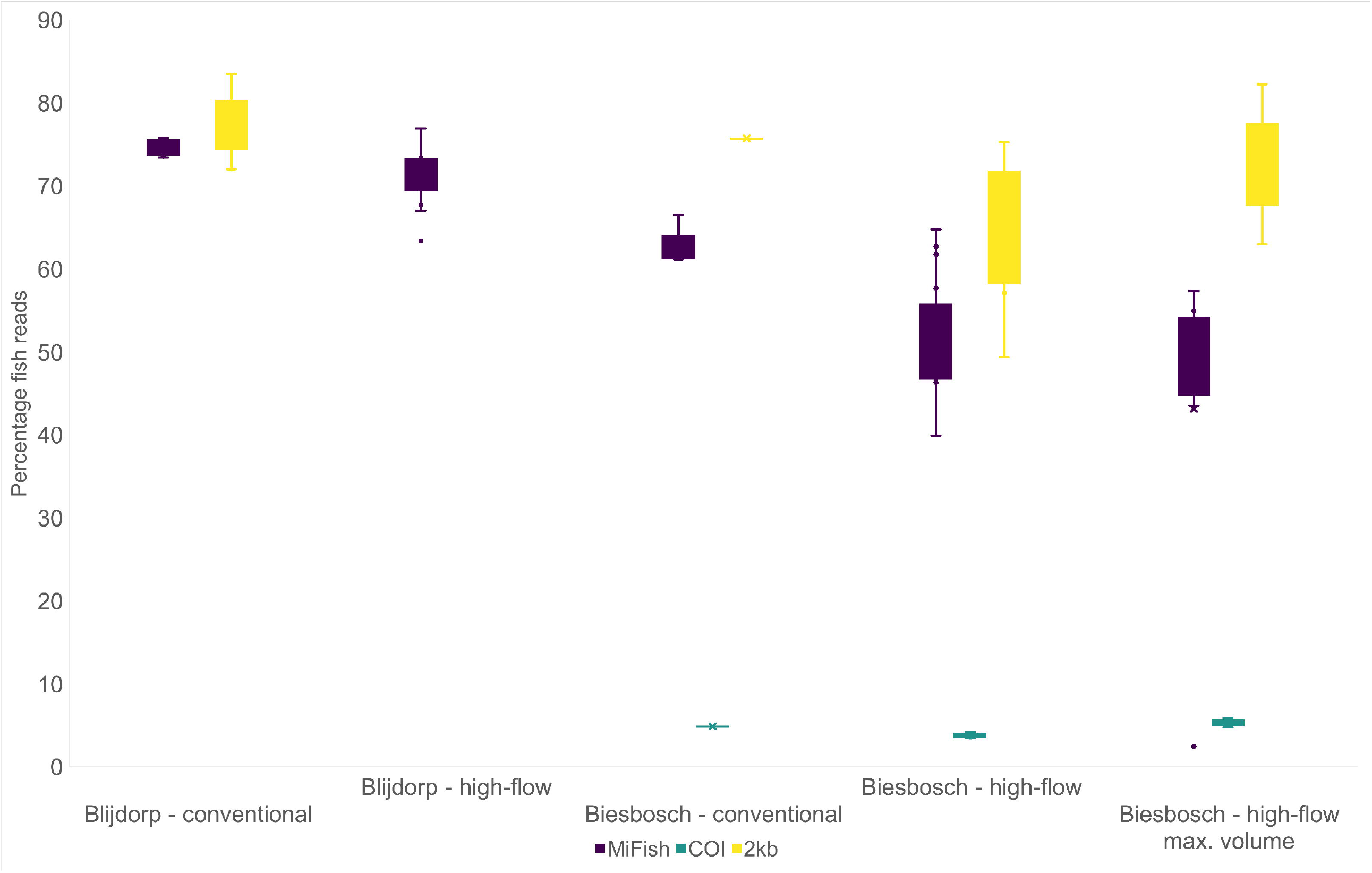
Percentage of metabarcoding reads classified as fish in two location using two filter types; Percentage of reads classified as fish (Actinopterygii & Elasmobranchii), from metabarcoding sequencing data obtained from eDNA water samples from a zoo aquarium (Blijdorp) and during a sturgeon release (Biesbosch) using a multi-marker approach (MiFish, COI & 2kb) and using two filter types: conventional 1.2 μm disk filter (2L) and a low cost cigarette filter (2L + max volume (8.3l)).

With metagenomics, 769,546 reads were identified as fish, covering 0.50% to 5.83% of total reads within an eDNA sample (Supplementary Table 4). In Blijdorp eDNA, more reads were identified as fish in the high-flow filters (4.51% ± 1.08, mean ± SD) compared to the conventional filters (1.51% ± 0.79, mean ± SD) (Fig. 4). Remarkably, the opposite was found in the Biesbosch, where 0.96% ± 0.46 (mean ± SD) of reads were identified as fish with the high-flow filters (Fig. 4) compared to 2.38 % ± 0.80 (mean ± SD) observed with the conventional filters. The high-flow filter used to its maximum capacity (8.3 L) in the Biesbosch showed a higher fish ratio (1.57%) compared to the high-flow filters where only 2 liters of water were filtered (mean = 0.96%). The Biesbosch Bucket eDNA samples yielded approximately between 4.51% (conventional filter) and 5.83% fish reads (Supplementary Table 4).

**Figure 4.**
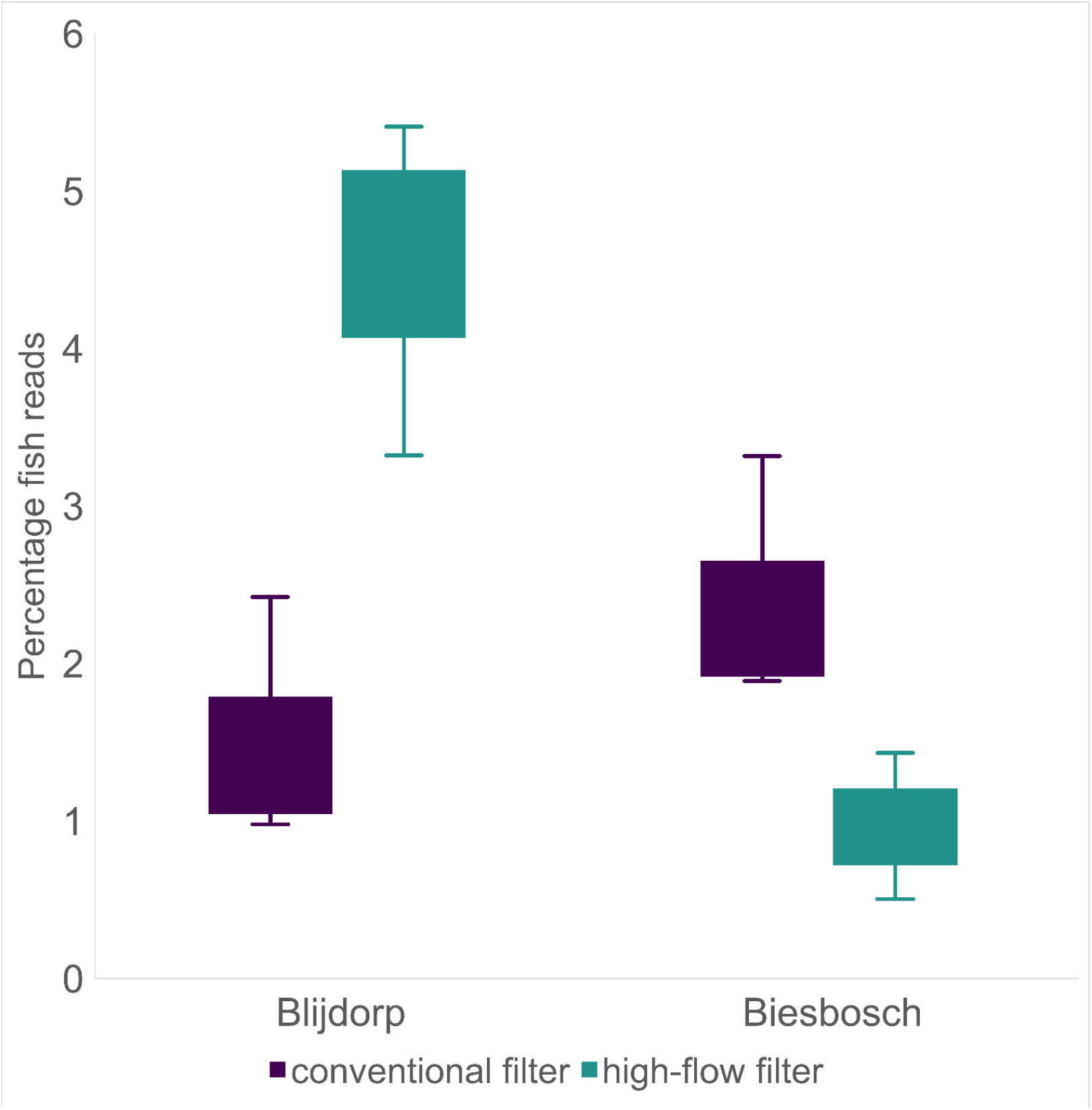
Percentage of metagenomic reads classified as fish in two location using two filter types; Percentage of reads classified as fish from metagenomic sequencing data obtained from eDNA water samples collected at a zoo aquarium (Blijdorp) and sturgeon release (Biesbosch) using two filter types: conventional 1.2 μm disk filter and a high-flow cigarette filter.

### Methylation patterns

Across all locations and filter types, 123, 32 and 124 methylated sites were detected for 5mC, 5mCG and 6mA on the 1761 bp 18S region, respectively. The coverage differed between methylation types and (pooled) samples with an average sequencing coverage of methylated sites of 38.9 ± 51.0 (mean ± SD) for 5mC, 37.0 ± 49.3 (mean ± SD) for 5mCG and 39.1 ± 50.8 (mean ± SD) for 6mA. For all three methylation types, Blijdorp samples taken with a conventional filter only had sites with 3 times coverage or less. Biesbosch Bucket high-flow filter had the highest coverage with over 150 times coverage for each dataset. Overall, the average rate of methylation per site for 5mC, 5mCG and 6mA were 21.0 ± 30.1 (mean ± SD), 65.4 ± 13.4 (mean ± SD) and 0.51 ± 2.06 (mean ± SD) for, respectively. Methylation rates did not differ between samples for 5mC (Kruskal-Wallis, X^2^ = 137.62, df = 176, *p* = 0.9854), 5mCG (Kruskal-Wallis, X^2^ = 98.908, df = 104, *p* = 0.836) nor 6mA (Kruskal-Wallis, X^2^ = 53.264, df = 92, *p* = 0.9996). One site showed significantly different methylation patterns between pooled samples in 5mC as well as 5mCG methylation (AY544133.1:1717), while two methylated sites were identified as significantly different between pooled samples in 6mA methylation (AY544133.1:198 and AY544133.1:1677). Following pairwise comparisons of all eDNA samples, some showed significantly differentiated methylated sites, but these did not overlap with the three sites that were determined overall significantly different.

## Discussion

### Species composition controlled environment

Based on the species present in the Blijdorp aquarium, fish species diversity was higher than expected. A total of fifteen fish species were detected in Blijdorp eDNA using metabarcoding, including all seven species that were present in the aquarium during the time of water collection: *Acipenser sturio, Dasyatis tortonesei, Labrus mixtus, Mustelus asterias, Raja brachyura, Scomber scombrus* and *Sparus aurata*. Some additional species that were detected with metabarcoding are part of the fishes’ diet (e.g. *Osmerus eperlanus*), while other species were present elsewhere in the zoo (e.g. *Carcharhinus* sp. *& Triakis semifasciata*). The eDNA of these species located elsewhere was likely pumped between aquaria through the zoo’s filtration-/refreshing system or transferred with tools or nets.

In metagenomics, eight fish species were missing compared to the metabarcoding dataset: *Atherina boyeri, Carcharhinus* sp., *Clupea pallasii, Dicentrarchus labrax, Gymnothorax* sp., *Osmerus eperlanus, Osphronemus goramy & Triakis semifasciata*. The lack of these species in metagenomic sequencing could be due to several factors, including 1) the lack of reference genomes limiting detection rates and 2) the extreme sensitivity of metabarcoding picking up rare eDNA. The lack of reference genomes was apparent in two species (*Osphronemus goramy & Triakis semifasciata*) which had no publicly available reference genome, severely limiting the detection possibilities. Two other species were only identified to genus level by metabarcoding (*Carcharhinus* sp. & *Gymnothorax* sp.). At the time of water collection, *Carcharhinus acronotus, Carcharhinus limbatus* and *Carcharhinus plumbeus* were present elsewhere in the zoo, but all three species had no reference genome available. Similarly, and *Gymnothorax favagineus, Gymnothorax javanicus* and *Gymnothorax isingteena* were present elsewhere in the zoo, but only *G. javanicus* had a reference genome available. Moreover, *Sprattus sprattus*, which was detected with metagenomics, might have been wrongly assigned as *Clupea pallasii* in MiFish metabarcoding. The 172 bp amplicon of these two species are highly similar, with only 1 or 2 bp difference depending on exact haplotype, making faulty assignment with bioinformatics likely. As *Sprattus sprattus* was part of the diet at the time of eDNA collection, this is the most probable origin of this species. Three other species (*Atherina boyeri, Dicentrarchus labrax, Osmerus eperlanus*) were not present in the sturgeon aquarium, but located elsewhere in the zoo, emphasizing the sensitivity of PCR in metabarcoding compared to metagenomics. The eight species detected in Blijdorp eDNA metagenomics were all present in the aquarium where the water was collected, or they were part of the fishes’ diet. As metagenomics is less sensitive to small amounts of eDNA originating outside of the eDNA collection location, it could be a more suitable technique for local species detection and corresponding biodiversity estimates. However, as this comes at the cost of potentially missing rare species present at the collection location, complementary usage of metabarcoding and metagenomics gives the best insight in (local) species presence.

### Species composition field settings

More species were identified in Biesbosch metagenomic sequencing compared to metabarcoding eDNA. Only five fish species were identified with metabarcoding, while metagenomics detected 10 species. Being part of the major riverine systems in the Netherlands, at least several dozen fish species are present in the Biesbosch nature reserve (Vrooman et al. 2021). The lack of species in observed in the Biesbosch metabarcoding data might be caused by the high relative abundance of *Acipenser sturio* eDNA. The high abundance of one species may have caused uneven amplification during PCR, a process known as species masking, thereby underrepresenting or even omitting lesser abundant fish species (Evans et al. 2016). The proportion of *Acipenser sturio* eDNA compared to other fish eDNA was higher in Biesbosch eDNA than in Blijdorp eDNA (97.71%, and 91.13% respectively). While the proportion of *Acipenser sturio* compared to other fish species was high in both locations, the proportion in Biesbosch eDNA potentially crossed a threshold where species masking is more likely to occur. In Biesbosch eDNA samples, only one species (*Neogobius fluviliatis*) was observed in metabarcoding samples, but not in metagenomics samples. At the time of this study, this species did not have a reference genome publicly available, limiting the detection possibilities using metagenomics. Other *Neogobius* spp. were present in the Kraken2 and BLAST reference databases, but these did not lead to the detection of *N. fluviliatis*, highlighting the specificity of the pipeline used in the present study.

In this study, metagenomic sequencing costs per sample were 5 times higher (€48/sample) than metabarcoding, excluding costs for PCR workflow. However, in the Biesbosch samples, metagenomic sequencing was able to provide better insight into species occurrence due to the lack of species masking occurring during PCR. In addition, metagenomic data contains species information of the entire tree of life (ToL). For this study, only a fraction (5% or less) of all metagenomic sequencing data is used, as we focus on a specific taxonomic group. While ToL metabarcoding would require additional PCRs and sequencing for each taxonomic group, this information is already provided by metagenomic sequencing. Thus, in case of ToL eDNA studies, metagenomic sequencing might be more cost-efficient than metabarcoding, in addition to the reduced time spent on laboratory tasks.

### Relative read counts

The differing relative read counts of overlapping species in metabarcoding and metagenomics in Blijdorp eDNA suggest that PCR amplification is more effective for some species than for others. Some species were overrepresented in metabarcoding compared to metagenomics, while others were underrepresented. This uneven amplification is known as primer bias and is extensively covered in literature (Polz and Cavanaugh 1998, Elbrecht and Leese 2015, Piñol et al. 2015, Ficetola and Taberlet 2023, Serite et al. 2023). The differences in relative read counts between metabarcoding and metagenomics can be affected by other factors including, but not limited to, genome sizes, ratios between mitochondrial DNA (mtDNA) and nuclear DNA (ntDNA), and excretion rates of mtDNA and ntDNA. In addition, the 200-fold discrepancy between metabarcoding and metagenomics of *Dasyatis tortonesei* relative read counts highlighted the importance of a reference genome. The reference data of this species was limited to a mitochondrial genome, meaning that all BLAST hits were made on this limited portion of the genome. While the mitochondrial genome is highly abundant compared to the nuclear genome, and therefore often used for metabarcoding, its size is a fraction of the total genome size. Notably, of the physiologically similar *Raja brachyura*, none of the >3500 reads identified with metagenomics were identified as mitochondrial genome, indicating that *D. tortonesei* likely contributes highly to the eDNA, but is simply not detected using metagenomics. Taken together, our observations highlight the power and sensitivity of PCR and simultaneously show the importance of complete reference databases. Therefore, the translation of relative read counts to abundance and/or biomass should be made with great caution in metabarcoding as well as metagenomics.

### Species classification

The overall percentage of sequenced reads classified as fish species was 63.4% for MiFish and 7.94% for COI, which is similar to other eDNA studies conducted in European waters (Collins et al. 2019, Van Driessche et al. 2023). The classification rates from Blijdorp eDNA were generally higher than those from Biesbosch, likely due to the density of animals being higher in Blijdorp and the fact that eDNA is less dispersed in an aquarium compared to a river. In addition, it is likely that less other DNA sources (Bacteria, Fungi, algae, invertebrates, etc.) are present in the controlled Blijdorp setting, compared to the Biesbosch environment, increasing the ratio of fish DNA. The metagenomic fish classification rates of Biesbosch eDNA (0.50% to 5.83%) were comparable to other studies conducted in field settings, where anywhere between <0.000001% to approximately 24% have been reported as eukaryotic classification rate from aquatic systems (Cowart et al. 2018, Bell et al. 2023, Manu and Umapathy 2023, Lui and Nielsen 2024, Curto et al. 2025, Patin et al. 2025, Serivichyaswat et al. 2025). This large variance observed in literature likely originates in the various locations, filter types, eDNA extraction protocols, sequencing providers and bioinformatic pipelines used. While the classification rates of metagenomic eDNA could be increased by using more lenient filtering steps (e.g. decreasing the minimum percentage identity), thereby allowing for more intraspecific variation, additional tools would be necessary to verify the species identification (Curto et al. 2025).

### Alternative amplicons Acipenseridae spp

Alternative Acipenseridae spp. amplicons were observed in the metabarcoding data across all eDNA samples and across all three primer sets. The fragments were classified as Acipenseridae spp., but with low percentage identity scores of ∼97% (MiFish & 2kb) or even < 93% (COI). The MiFish and 2kb alternative amplicons had a similar relative read counts with 12.6% ± 11.7 (mean ± SD) and 13.7 % ± 11.1 (mean ± SD), respectively, and featured several indels and dozens of single nucleotide polymorphisms (SNPs) on the 12S and 16S rRNA. Alternative amplicons were also observed in all eDNA samples amplified with the COI primer set, with relative read counts of 41.4% ± 6.13 (mean ± SD) showing higher relative read counts and more consistency than MiFish and 2kb. Several SNPs were observed in the alternative COI amplicon, of which the majority was classified as missense mutation. As the alternative amplicons were present in Blijdorp, the Biesbosch Holding tank and the Biesbosch Bucket, the most likely origin of the alternative amplicons was *Acipenser sturio*.

To determine whether the origin of these fragments were nuclear mitochondrial DNA fragments (NUMTs) (Gellissen et al. 1983), all amplicons were mapped to the *Acipenser sturio* nuclear genome (GCA_977009215) using minimap2 (Li 2018). All amplicons mapped to the first scaffold of the assembly; MiFish amplicons mapped to HiC_scaffold_1: 72,413,745-72,413,910, 2kb amplicons mapped to: HiC_scaffold_1: 72,413,951-72,415,898 and the primary mappings of COI amplicons were in: HiC_scaffold_1: 72,430,181-72,430,484. In all three primer sets, the most common amplicon mapped perfectly to the mitochondrial genome, while the second most common amplicon (the alternative amplicon) mapped perfectly to HiC_scaffold_1. Therefore, the most likely origin of the alternative metabarcoding amplicons is NUMTs. While the region in HiC_scaffold_1 clearly represents a NUMT, the differing proportion of alternative amplicons between primer sets suggests that more NUMTs exist in the *Acipenser sturio* genome. In other fish species, NUMTs were found to be extremely common, with hundreds of NUMTs detected in some species (Antunes and Ramos 2005). While the occurrence of NUMTs in mtDNA metabarcoding is unsurprising, the extremely high abundance could impact the reliability of metabarcoding sequencing data (Turanov et al. 2024).

### Filter types

The high-flow filters did not increase the proportion of target (fish) eDNA captured in the Biesbosch field setting. However, the high-flow filters did improve capture of target DNA in the controlled Blijdorp environment. These findings are unexpected, as it was hypothesized that the higher flow rate and increased pore size would increase capture rates of larger clumps of multicellular organisms, while simultaneously passing smaller non-target particles. We expected that these potential benefits would become apparent in the murky waters of field setting, while the benefits would be redundant in the cleaner water of the controlled environment. The surprising result could partially be explained by the low filtration volume of the high-flow filter, as most of these filters were not used to their maximum capacity in this study, unlike the conventional filters. Other studies have shown that filtering until maximum capacity improves species recovery, consistency and ecosystem representation (Govindarajan et al. 2022, Nester et al. 2024). Similarly, the single high-flow filter which was used to its maximum filtering capacity in this study yielded higher rates of target species. However, since this study incorporated only a single filter, future repeated measures are needed to confirm the hypothesis. In addition to the benefits of increased capture rates of multicellular organisms, the high-flow filters and 3D printed filter holders are convenient to use during fieldwork on land or at sea and are very suitable for remote fieldwork due to their global availability and low cost. While other low-cost, globally available, filter solutions have been reported such as coffee filters or passive sponge filtrations (Chen et al. 2024, Mlinarec et al. 2025), the cigarette filter could improve handling and decrease costs.

More diverse fish communities were observed with metagenomics using the high-flow filter, compared to the conventional filter, in the controlled environment as well as the field setting, demonstrating the high-flow filter can outperform the conventional filter in terms of species detection. A similar pattern was observed in metabarcoding of the controlled environment in Blijdorp, where more species were detected using the high-flow filter. This pattern was not observed in the Biesbosch field setting, where a poor overlap in species was found between the two filter types. The extremely high abundance of *Acipenser sturio* eDNA copies in Biesbosch metabarcoding may have caused species masking in both filter types, leading to irregular and limited species detection (Evans et al. 2016). In addition to the additional species observed with metagenomics using the high-flow filters, the mean target fragment length observed in the high-flow filters was 30% to 60% higher than the conventional filter in the Blijdorp samples and Biesbosch Holding tank samples. Fragment length can be essential for species detection in metagenomics, as longer DNA fragments inevitably hold more information than short ones.

### Methylation patterns

Using metagenomic data from all sampling environments and filter types, it was possible to analyze epigenetic patterns of three methylation types: 5mC, 5mCG and 6mA. As the coverage between samples was highly variable, genome wide epigenetic analysis was not possible, but methylation calling and subsequent analysis of a 1761 bp section of the 18S gene revealed several differentially methylated nucleotides. While methylation of the 18S gene is not directly linked to any life-history traits, other studies have linked methylated eDNA to age estimations and stress-responses of aquatic organisms (Crossman et al. 2021, Zhao et al. 2023, Ruiz et al. 2025). Using Oxford Nanopore’s native sequencing allow analyses of species diversity as well as epigenetic patterns from the same sequencing run. Where most biodiversity-orientated studies would have additional expenses of bisulfite sequencing to provide epigenetic information, Oxford Nanopore’s native sequencing can always provide methylation patterns from a sequencing run, even retroactively. We strongly recommend future eDNA studies to take the epigenetic analyses in consideration, as to obtain a more complete picture of local biodiversity (species diversity, genetic diversity, epigenetic patterns) from a single eDNA sample (Valk et al. in prep).

## Conclusions

While metabarcoding excels in species detection, it might overestimate the number of local species present due to its sensitivity to rare (foreign) DNA. Metagenomics can be used as complementary technique to circumvent the inherent primer bias and potential species masking in metabarcoding. In addition, Oxford Nanopore’s native sequencing, allows insight in epigenetic patterns. The high-flow, low-cost, cigarette filter and 3D printed filter holder presented in this study allow affordable eDNA filtration, while often obtaining superior insights into fish communities compared to the conventional 1.2 μm disk filter.

## Acknowledgements

We thank Diergaarde Blijdorp/Rotterdam Zoo and their employees for their assistance on the water collection of the sturgeon aquarium. In addition, we thank MIGADO, Sportvisserij Nederland, ARK Rewilding the Netherlands and WWF-NL, who facilitated the sturgeon release in the Biesbosch and assisted in water collection. Lastly, we thank Vittorio Saggiomo for his contribution to the design of the 3D-printed filter holders.

## Data availability

Data and code are available on https://github.com/DvBMolEc/ONT_native_kraken2_BLAST. All sequence data are available at https://www.ebi.ac.uk/ena/browser/view/PRJEB96019 (BioProject PRJEB96019).

